# Revealing the Complexities of Metabarcoding with a Diverse Arthropod Mock Community

**DOI:** 10.1101/433607

**Authors:** Thomas W. A. Braukmann, Natalia V. Ivanova, Sean W. J. Prosser, Vasco Elbrecht, Dirk Steinke, Sujeevan Ratnasingham, Jeremy R. deWaard, Jayme E. Sones, Evgeny V. Zakharov, Paul D. N. Hebert

**Affiliations:** Centre for Biodiversity Genomics, University of Guelph, Guelph ON, Canada N1G 2W1; Department of Integrative Biology, University of Guelph, Guelph ON, Canada N1G 2W1; School of Environmental Sciences, Uinversity of Guelph, Guelph, ON, Canada N1G 2W1

## Abstract

DNA metabarcoding is an attractive approach for monitoring biodiversity. However, it is subject to biases that often impede detection of all species in a sample. In particular, the proportion of sequences recovered from each species depends on its biomass, mitome copy number, and primer set employed for PCR. To examine these variables, we constructed a mock community of terrestrial arthropods comprised of 374 BINs, a species proxy. We used this community to examine how species recovery was impacted when amplicon pools were constructed in four ways. The first two protocols involved the construction of bulk DNA extracts from different body partitions (<u>Bulk Abdomen, Bulk Leg</u>). The other protocols involved the production of DNA extracts from single legs which were then merged prior to PCR (<u>Composite Leg)</u> or PCR-amplified separately (<u>Single Leg</u>) and then pooled. The amplicon generated by these four treatments were then sequenced on three platforms (Illumina MiSeq, Ion Torrent PGM and Ion Torrent S5). The choice of sequencing platform did not substantially influence species recovery, other variables did. As expected, the best recovery was obtained from the <u>Single Leg</u> treatment, but the <u>Bulk Abdomen</u> produced a more uniform read abundance than the <u>Bulk Leg</u> or <u>Composite Leg</u> samples. Primer choice also influenced species recovery. Our results reveal how variation in protocols can have substantive impacts on perceived diversity unless sequencing coverage is sufficient to reach an asymptote. Although metabarcoding is a powerful approach, further optimization of analytical protocols is crucial to obtain reproducible results and increase its cost-effectiveness.

## Introduction

It is generally accepted that we have entered a period of unprecedented global biodiversity loss (Pimm et al. 2014, Vogel 2017). Halting it will require the capacity to quantify shifts in species composition rapidly and on a far larger scale than ever before so we can better understand and manage ecosystems (Ji et al. 2013; Cristescu 2014; Moriniere et al. 2016, Waldron et al 2017). As arthropods account for the majority of terrestrial biodiversity (Medeiros et al. 2013), they are an obvious target for bio-surveillance. Although they are easily collected in large numbers (Russo et al. 2011), the subsequent processing and identification of specimens has traditionally been a barrier to large-scale monitoring programs (Bassett et al. 2012). DNA barcoding, the use of short standardized gene regions to discriminate species, breaks this barrier by enabling non-taxonomists to identify specimens once a reference sequence library is established (Hebert et al. 2003; Hebert and Gregory 2005).

DNA barcode studies initially focused on developing the analytical protocols required to construct a specimen-based reference library (Hebert et al. 2003; Hebert et al. 2004). Although improved protocols have reduced costs, making it possible to analyze millions of single specimens (Hajibabaei et al. 2005; Ivanova et al. 2006; Hebert et al. 2018), they are still too expensive to support large-scale bio-monitoring programs. However, by coupling a DNA barcode reference library with the analytical capacity of high-throughput sequencers (HTS), DNA metabarcoding provides a path to rapid, low-cost assessments of species composition (Hajibabaei et al. 2011; Yu et al. 2012; Brandon-Mong et al. 2015; Moriniere et al. 2016). It achieves this goal by the amplification and sequence characterization of the barcode region from bulk DNA extracts which can then be assigned to operational taxonomic units (OTUs) that can be queried against reference sequences to ascertain their source species (see Cristescu 2014). Studies have now employed this approach to assess species composition in communities of aquatic and terrestrial arthropods (Ji et al. 2013; Beng et al. 2016; Elbrecht et al. 2017B), vertebrates (Sato et al. 2017), diatoms (Vasselon et al. 2017), and fungi (Bellemain et al. 2012; Aas et al. 2017; Tedersoo et al. 2018). Metabarcoding can reveal more species than morphological approaches while requiring far less time (Ji et al. 2013; Yu et al. 2012; Vivien et al. 2016; Brandon-Mong et al. 2015; Elbrecht et al. 2017A; Hebert et al. 2018; Shokralla et al. 2015; Elbrecht et al. 2017B).

Despite the advantages of metabarcoding, several factors often complicate the recovery of all species in a sample. Firstly, DNA templates derived from the species in a mixed sample are often differentially amplified (Elbrecht and Leese 2015; Pinol et al. 2015; Elbrecht and Leese 2017; Tedersoo et al. 2018). Such bias can arise from either the DNA polymerase (Nichols et al. 2018; Pan et al. 2014, Dabney and Meyer 2012) or the primers (Clarke et al. 2014) employed for PCR. Polymerase bias involves the differential amplification of templates as a result of their variation in sequence motifs, GC content, or length (Nichols et al. 2018; Pan et al. 2014, Dabney and Meyer 2012). Primer bias is due to either varying levels of primer mismatch or template degradation (Clarke et al. 2014; Elbrecht and Leese 2015). The impact of primer mismatches can often be reduced by either lowering annealing temperatures or by raising the degeneracy of the primers (Clarke et al. 2014; Elbrecht and Leese 2017). However, these ‘solutions’ have a downside; they often increase the amplification of non-target sequences such as bacterial endosymbionts or mitochondrial pseudogenes, which is especially problematic for eDNA samples (Smith et al. 2012; Song et al. 2008; Macher et al. 2018).

The capacity of metabarcoding to recover all species in a bulk sample is further complicated because the component species typically vary by several orders of magnitude in mass and hence in copy numbers of the target template. Unless other factors intervene, this variation in template number means that large-bodied species are more likely to be recovered (Brandon-Mong et al. 2015). Because of this effect (in addition to primer bias), efforts to infer species abundance from the read counts obtained in metabarcoding studies are at best weak (Elbrecht and Leese 2015; Pinol et al. 2014). Correction factors can improve such estimates (Thomas et al. 2015; Vasselon et al 2017), but any method based on the analysis of bulk DNA extracts will fail to accurately estimate species abundance.

In addition to factors complicating the recovery of sequences from all species in a bulk sample, sequence variation introduced during PCR, library preparation, and sequencing can make it difficult to assign sequences to their source species (Tedersoo et al. 2018). PCR error can be reduced by the use of high-fidelity polymerases (Lee et al. 2016; Potapov et al. 2017), but it is more difficult to escape complexities introduced by sequencing error because all second-generation sequencers have error rates (e.g. 1–2%) that are high enough to complicate the discrimination of closely-related species. Third-generation sequencers, such as Pacific Biosciences Sequel (e.g. Hebert et al. 2018), can produce much lower error rates, but they currently generate too few reads (circa 0.2 million/run) to reveal all species in a taxonomically diverse sample (Tedersoo et al. 2018). As a consequence, despite their high error rates, second-generation platforms (Illumina, Ion Torrent) are commonly used for metabarcoding as they produce many millions of reads per run (Mardis et al. 2013, Cristescu 2014). Illumina sequencers generate more reads (20–250 million/run) with lower error rates than Ion Torrent platforms, but the latter instruments can deliver longer reads more rapidly (Mardis et al. 2013). It is unclear how severely the choice of HTS platform affects species recovery as their performance has rarely been compared in eukaryotes (Divolli et al. 2018). However, work on microbial communities found general agreement between platforms although Ion Torrent reads were lower quality and more length variable than those from Illumina (Salipante et al. 2014; Tessler et al. 2017).

To examine factors influencing the success in recovering species through metabarcode analysis, we assembled a mock community that included a single representative of 374 species. We subsequently used this community to examine the impacts of DNA source, extraction method, PCR protocol, target template, and sequencing platform on species recovery. Specifically, we compared results obtained by analyzing four amplicon pools on three sequencing platforms (Illumina MiSeq, Ion Torrent PGM, Ion Torrent S5). Two of these amplicon pools derived from the PCR of bulk DNA extracts (abdomen, leg) to test the impact of tissue type. Two other amplicon pools derived from DNA extracts of single legs that were analyzed by pooling prior to or after PCR. Finally, we examined species recovery for two amplicons of differing length on the S5. The overall analytical approach involved evaluation of the relationship between read depth and species recovery for these treatment variables.

## Material and Methods

### Assembly of Mock Community

We began the assembly of a mock community by obtaining COI sequences from 3,044 insects collected in Malaise traps deployed near Cambridge, Ontario, Canada. A DNA extract was prepared from a single leg from each specimen employing a membrane-based protocol (Ivanova et al. 2006). The 658 bp barcode region of COI was amplified and then Sanger sequenced, to link a haplotype to each individual specimen. Amplicons were generated using a primer cocktail of C_LepFolF and C_LepFolR (Hernández-Triana LM et al. 2014) with initial denaturation at 94 °C for 2 min followed by 5 cycles of denaturation for 40 s at 94 °C, annealing for 40 s at 45 °C and extension for 1 min at 72 °C; then 35 cycles of denaturation for 40 s at 94 °C with annealing for 40 s at 51 °C and extension for 1 min at 72 °C; and a final extension for 5 min at 72 °C (Ivanova et al. 2006; Hebert et al. 2018). Most reactions generated a 709 bp amplicon comprised of 658 bp of COI plus 51 bp of forward and reverse primers. A few amplicons were slightly shorter as a result of deletions in the COI gene. Unpurified PCR products were diluted 1:4 with ddH_2_O before 2 μl was used as the template for a cycle sequencing reaction. All products were sequenced following standard procedures on an ABI 3730×1 DNA Analyzer (Applied Biosystems, Foster City, California, USA).

Because some specimens could not be identified to a species level, we employed the Barcode Index Number (BIN) system which examines patterns of sequence variation at COI to assign each specimen to a persistent species proxy (Ratnasingham and Hebert 2013). The overall analysis provided sequence records for 803 BINs. From this total, we selected 374 BINs that met two criteria. They showed >2% COI sequence divergence from their nearest-neighbor, and they were taxonomically distant. The resulting mock community included representatives of 10 orders and 104 insect families. Supplemental Table 1 provides a list of the taxa included in the mock community as well as details on vouchers, their body size (as estimated by abdominal mass), and the GC content of their COI. Following selection of the specimens for inclusion in the mock community, fresh DNA extracts were made following the four protocols described below, and the amplicon pools generated from them were subsequently analyzed on three sequencing platforms.

**Table 1.**
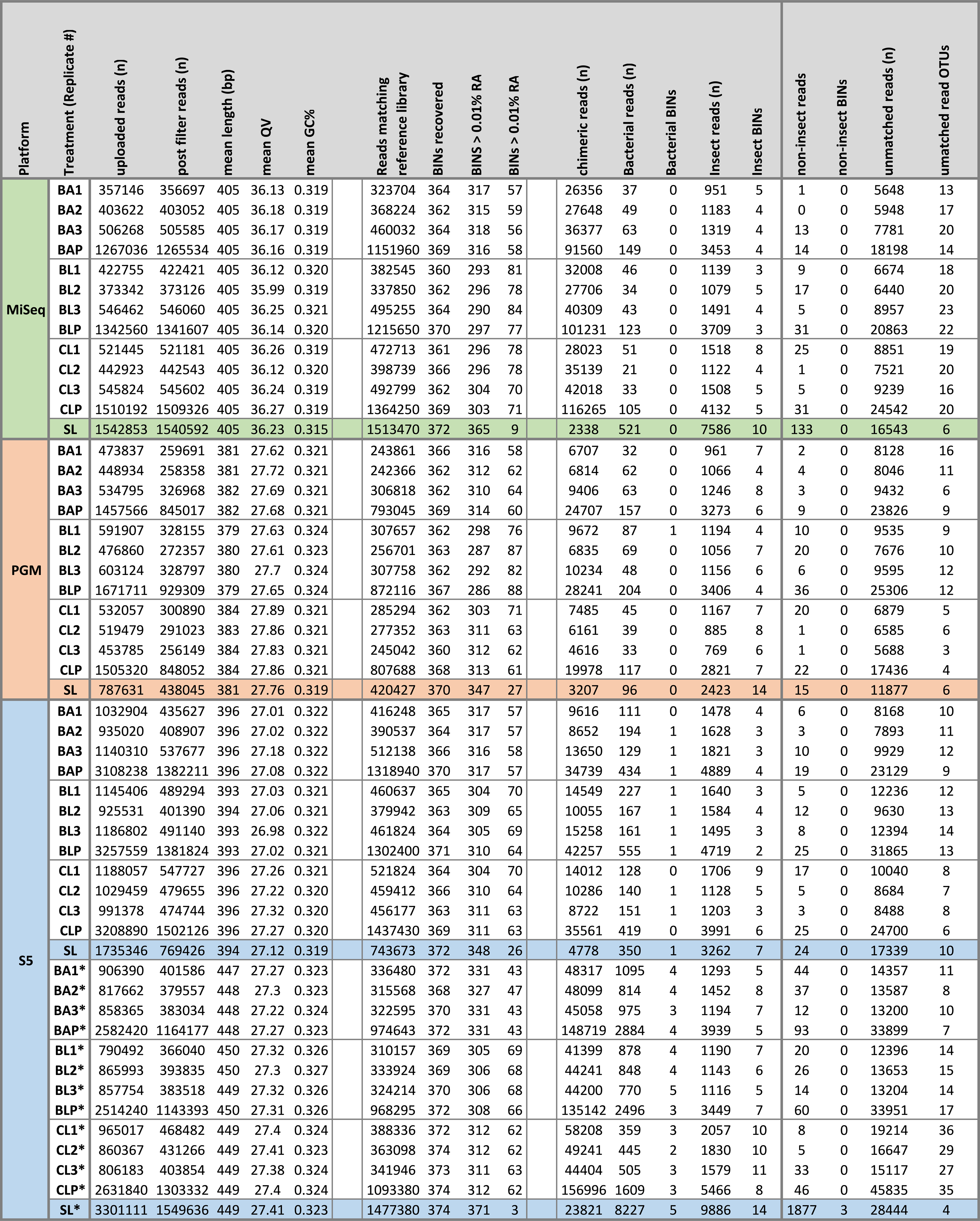
Summary of run results for all treatments. mBRAVE filtering and BIN recovery including false positives are indicated for the four amplicon pools (BA = <u>Bulk Abdomen,</u> BL = <u>Bulk Leg,</u> CL = <u>Composite Leg,</u> SL = <u>Single Leg</u>). Replicates are numbered 1–3 and pooled replicates are denoted by a P. BINs (Barcode Index Number) for not reference library matches were only counted if their relative abundance was greater than 0.01%. All results are based on the analysis of a 407 bp amplicon except those marked with a * which are based on a 463 bp amplicon.

### Experimental design for metabarcode analysis

Species recovery was compared for amplicon pools generated by four protocols (Figure 1). Two involved the analysis of amplicons generated from bulk DNA extracts derived from two tissues (<u>Bulk Abdomen, Bulk Leg)</u>. The other two treatments involved the initial extraction of DNA from individual legs. The resultant DNA extracts were either pooled prior to PCR to create the <u>Composite Leg</u> treatment or separately amplified and subsequently pooled to create the <u>Single Leg</u> treatment (Figure 1). Although the initial design called for the same specimens to be included in each mock community, this was not possible. The <u>Composite Leg</u> and <u>Single Leg</u> treatments did include the selected array of 374 BINs. However, five of their source specimens either lacked an abdomen or another leg for inclusion in the <u>Bulk Abdomen</u> or <u>Bulk Leg</u> treatments. As a result, five BINs, generally belonging to the same order as the excluded ones, were employed as replacements to maintain 374 BINs per treatment (BOLD:AAA2323, BOLD:AAA2632, BOLD:AAF4234, BOLD:AAP6354; BOLD:ABV1240). Further details on the treatments are available at the following DOIs: <u>Bulk Abdomen</u> and <u>Bulk Leg</u>: dx.doi.org/10.5883/DS-NGS375A: <u>Composite Leg</u> and <u>Single Leg</u>: dx.doi.org/10.5883/DS-NGS375B.

**Figure 1.**
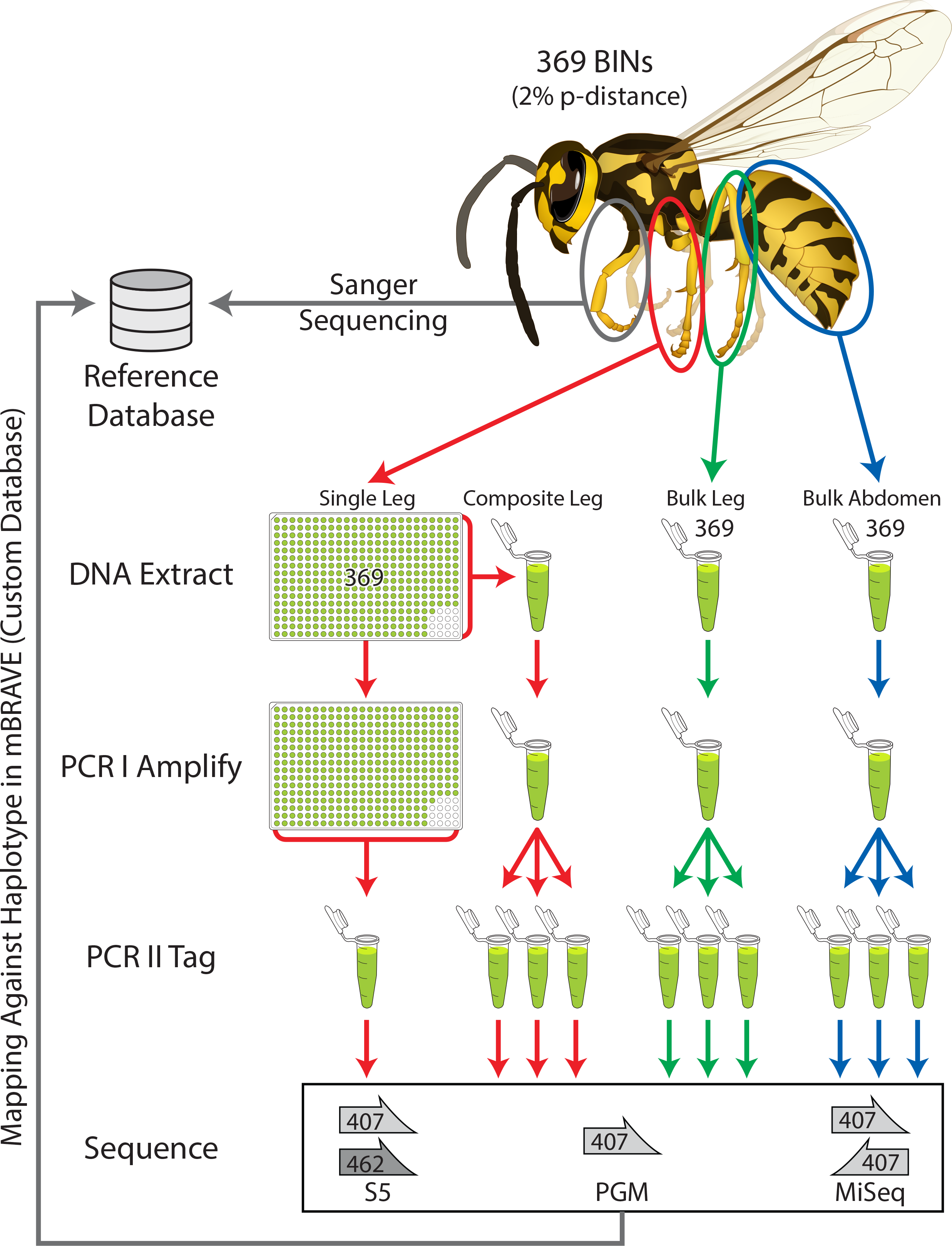
Protocol employed to examine species recovery from the mock community. Four amplicon pools were examined. Two derived from bulk DNA extracts (<u>Bulk Abdomen, Bulk</u> Leg). The others derived from DNA extracts from single legs that were either pooled (<u>Composite Leg</u>) or kept separate (<u>Single Leg)</u> prior to PCR. All four amplicon pools were sequenced on three platforms (Illumina MiSeq, Ion Torrent S5, and Ion Torrent PGM). There were three technical replicates for each treatment except <u>Single Leg</u>.

### Bulk DNA extractions and PCR

DNA extracts for the two bulk samples (<u>Bulk Abdomen, Bulk Leg)</u> were generated with a modified membrane-based protocol (Ivanova et al. 2006). Specifically, the bulk abdomens (combined mass = 1,062.8 mg) and bulk legs (combined mass = 30.9 mg) were lysed overnight in the same relative volume of insect lysis buffer (51.6 ml and 1.5 ml respectively), with 10 mg/ml of Proteinase K (Invitrogen). Following lysis, a 100 μl aliquot of each extract was mixed with 200 μl of binding mix and transferred to an EconoSpin^®^ column (Epoch Life Sciences) before centrifugation at 5000 g for 2 min. The DNA extracts were then purified with three wash steps. The first wash employed 300 μl of protein wash buffer before centrifugation at 5000 g for 2 min. Columns were then washed twice with 600 μl of wash buffer before being centrifuged at 5000 g for 4 minutes. Columns were transferred to clean tubes and spun dry at 10000 g for 4 min to remove any leftover buffer, then transferred to clean collection tubes and incubated for 30 min at 56°C to dry the membrane. DNA was subsequently eluted by adding 50 μl of 10 mM Tris-HCl pH 8.0 followed by centrifugation at 10000 g for 5 min. All DNA extracts were normalized to 3 ng/μl prior to PCR. All PCR reactions were composed of 5% trehalose (Fluka Analytical), 1× Platinum Taq reaction buffer (Invitrogen), 2.5 mM MgCl_2_ (Invitrogen), 0.1 μM of each primer (Integrated DNA Technologies), 50 μM of each dNTP (KAPA Biosystems), 0.15 units of Platinum Taq (Invitrogen), 1 μl of template, and HyClone^®^ ultra-pure water (Thermo Scientific) for a final volume of 6 μl.

### Construction of HTS libraries

Two rounds of PCR were used to generate the amplicon libraries destined for sequence characterization on the three platforms. Most first-round reactions employed a primer cocktail targeting a 407 and 421 bp region of COI and will be referred to as the 407 bp amplicon throughout the manuscript. The 407 bp region was amplified using MLepF1 (Hebert et al. 2004) and the 421 bp region with RonMWasp (Smith et al. 2012) as forward primers and LepR1 (Hebert et al. 2004) and HCO2198 (Folmer et al. 1994) as reverse primers (Table S2). An alternate first-round PCR targeted a 463 bp amplicon of COI; it was generated with another forward primer – AncientLepF3 (Prosser et al. 2016) (Table S2). In addition, the primers employed to generate amplicons for MiSeq analysis contained a Nextera transposase adapter (Table S2). All first-round PCRs were run under the same conditions with initial denaturation of 94 °C for 2 min, followed by 20 cycles of denaturation at 94°C for 40 s, annealing at 51°C for 1 min and extension at 72 °C for 1 min, with a final extension at 72°C of 5 min. Three technical PCR replicates were generated for three of the treatments – <u>Bulk Abdomen, Bulk Leg,</u> and <u>Composite Leg</u>.

Prior to the second PCR, first-round products were diluted 2x with dd H_2_0. Fusion primers were used to attach platform-specific unique molecular identifiers (UMIs) along with sequencing adaptors for Ion Torrent libraries and a flow cell bind for the MiSeq libraries (Table S2). The second PCR was run under the same conditions as the first round for reactions slated for analysis on the Ion Torrent platforms, but the samples for Illumina were amplified following manufacturer’s specifications with initial denaturation at 94°C for 2 min, then 20 cycles of denaturation at 94°C for 40 with annealing at 61°C for 1 min and extension at 72 °C for 1 min, followed by a final extension at 72°C of 5 min. Supplementary Table S2 provides all primer sequences and details on samples indexing.

For each platform, the UMI-labelled reaction products were pooled prior to sequencing. The sequence libraries for the S5 were prepared on an Ion Chef™ (Thermo Fisher Scientific) following manufacturer’s instructions while those for the PGM were prepared using the Ion PGM™ Hi-Q™ View OT2 400 Kit and the Ion PGM™ Hi-Q™ Sequencing Kit (Thermo Fisher Scientific). The PGM libraries were sequenced on a 318 v2 chip while the S5 libraries were sequenced on a 530 chip at the Canadian Centre for DNA Barcoding. Illumina libraries were sequenced (paired end) using the 300 bp reagent kit v3 on an Illumina MiSeq in the Genomics Facility of the Advanced Analysis Centre at the University of Guelph.

### Bioinformatics and analysis

Prior to uploading MiSeq runs, read libraries were paired using the QIIME (Caporaso et al. 2010) pair join script (join_paired_ends.py) with a minimum overlap of 20 bp and a maximum difference of 10%. All read libraries were uploaded to mBRAVE (http://mbrave.net/), an online platform for analyzing and visualizing metabarcoding data. The Quality Value (QV) of each sequence was evaluated and all records failing to meet any one of three quality standards were discarded: 1) mean QV<20; 2) >25% of bp with QV<20; 3) >5% of bp with QV<10. All reads were trimmed to be either 407 bp or 463 bp. Retained sequences were viewed as a match to a BIN in the custom Sanger reference library if their distance was <3% to any reference. All reads not matching a reference sequence were clustered at an OTU threshold of 2%. These OTUs were then queried against four other system libraries (insects, non-insect arthropods, non-arthropod invertebrates, bacteria). Standard analytical parameters were used for all treatments and sequencing platforms. All raw data is available in NCBI’s Short Read Archive (SRP158933). The three replicates for the <u>Bulk Abdomen, Bulk Leg,</u> and <u>Composite Leg</u> treatments were pooled.

OTU tables for each run were merged in R ver 3.4.4 (R Core Team 2018). To compare BIN accumulation across all samples, we randomly subsampled each run at different read depths for 10,000 replicates using a custom script (Supplemental material). To measure the BIN accumulation for each treatment, we compared the slopes between sequential points at eight read counts (10^2^, 10^3^, 10^3.5^ 10^4^, 10^4.5^, 10^5^, 10^5.5^,10^6^). Sequential points with a slope of less than 0.01 were viewed as indicating that an asymptote had been achieved.

To compare the different treatments and sequencing platforms, we reduced the data set to the 369 shared BINs. Read distributions were visualized using the JAMP v0.44 package (https://github.com/VascoElbrecht/JAMP) in R to produce a heat map using the “OTU_heatmap” function. Read distributions across BINs were compared using density graphs generated with ggplot2 v2.2.1 (Wickham 2009). The relative abundances of all BINs comprising greater than 0.01% of the overall reads were used to estimate Simpson’s index, Pielou’s mean evenness, and Renyi’s entropy implemented in the R package vegan v2.5-1 (Oksanen et al. 2018). Compositional dissimilarity between replicates and treatments were examined using a dendrogram based on the Bray-Curtis index and calculated with vegan. The values for the Bray-Curtis index were also used to generate a non-metric multidimensional scaling (NMDS) with vegan.

The relationships between read counts and body size, as measured by abdominal mass, and the read count and GC content of the COI amplicon were examined using Kendall Tau correlations in R ver 3.4.4 (R Core Team 2018). An analysis of similarity (ANOSIM) with 999 permutations was used to compare species recovery among treatment types, and sequencing platforms and between the two amplicons with the R package vegan v2.5-1 (Oksanen et al. 2018). All custom scripts are available as supplementary materials.

The relationships between the read count for each BIN and primer mismatches were investigated for the 407 bp and 463 bp amplicons. The number of mismatches were quantified by counting the number of nucleotide substitutions between the primer sequence and the template DNA for each BIN. Information on the DNA sequence for the forward primer binding sites was available from the Sanger reads for all 369 BINs. Calculation of mismatches was straightforward for the 463 bp amplicon as it involved a single forward primer. As the 407 bp amplicon was generated with two different forward primers, total mismatches were quantified based upon the forward primer with the best match to the template for each BIN. The same two reverse primers were employed to generate the 407 bp and 463 bp amplicons, requiring a similar approach, but with the complication that DNA sequence information for template DNA was not available from the Sanger sequence (as it was based on amplicons generated with the same reverse primers). As a result, a new reverse primer was employed to extend the sequence in a 3’ direction, an approach which delivered the desired sequence information for 203 of the 369 BINs. As a consequence, it was possible to examine the relationship between read counts and the number of mismatches between template and forward primer for all 369 BINs and for the total mismatch count for the forward and reverse primers for the 203 BINs with template sequences for both regions.

## Results

### Run quality

We first compared the output and quality of the reads from the HTS platforms. The S5 and MiSeq generated a similar number of reads (~ 1 million per replicate), while the PGM generated substantially fewer. About 60–65% of the MiSeq reads were filtered during merging of the paired-end reads, but subsequent filtering was minimal (< 1%). The PGM and S5 encountered a similar loss of reads as 45–50% of the raw reads were filtered (Table 1). The MiSeq reads showed more length consistency and higher quality than those from both Ion platforms, reflecting their near consistent QV versus the decline towards the 3’ end of the PGM and S5 reads (Figure S1).

### Read depth

Rarefaction curves were calculated for each of the four treatments and their technical replicates to ascertain if read depths were sufficient to recover all BINs (Figure 2). Although BIN recovery was high in all cases, the <u>Single Leg</u> treatment reached it with far fewer reads of the 407 bp amplicon than the other treatments (10^4–4.5^ versus10^4.5–5^ –Table S3). There was evidence of variation among platforms as the PGM needed more reads to achieve an asymptote than the S5 or MiSeq. BIN accumulation curves for the other treatments were similar, but the <u>Bulk Abdomen</u> showed a small, but consistent outperformance of the <u>Bulk Leg</u> and <u>Composite Leg</u> treatments. The target amplicon also had a substantial impact as just 10^3.5^ reads of the 463 bp amplicon were required for the <u>Single Leg</u> treatment to reach its asymptote (Table S3). The technical replicates showed little divergence on all platforms; they had similar BIN recovery, similar mean read counts per BIN, and similar coefficients of variation (Table S1). Pielou’s evenness, Simpson’s Index, Inverse Simpson’s Index, Renyi’s diversity, and Shannon Indices were also similar across treatments on all platforms (Table 2; Figure S2). Finally, density plots were similar among technical replicates for all treatments and platforms suggesting that different HTS platforms produced similar results for the different treatments (Figure 1; Figure S3).

**Figure 2.**
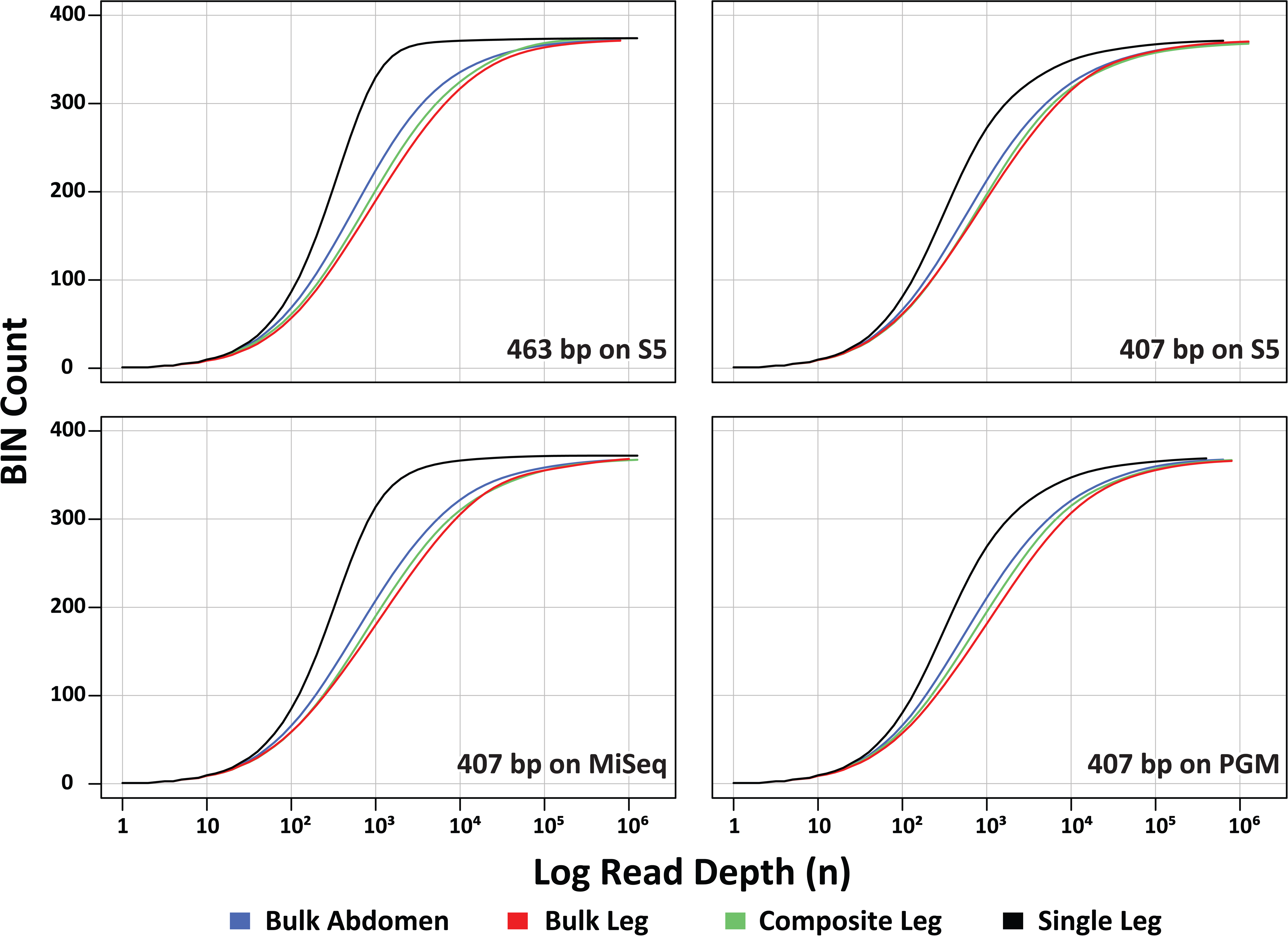
Rarefaction curves showing BIN recovery versus the number of sequences analyzed for the four amplicon pools (<u>Bulk Abdomen, Bulk Leg, Composite Leg, Single Leg</u>) on the three sequencing platforms. Two amplicon lengths (407 bp, 463 bp) were analyzed on the S5, but just one (407 bp) on the other platforms.

**Table 2.**
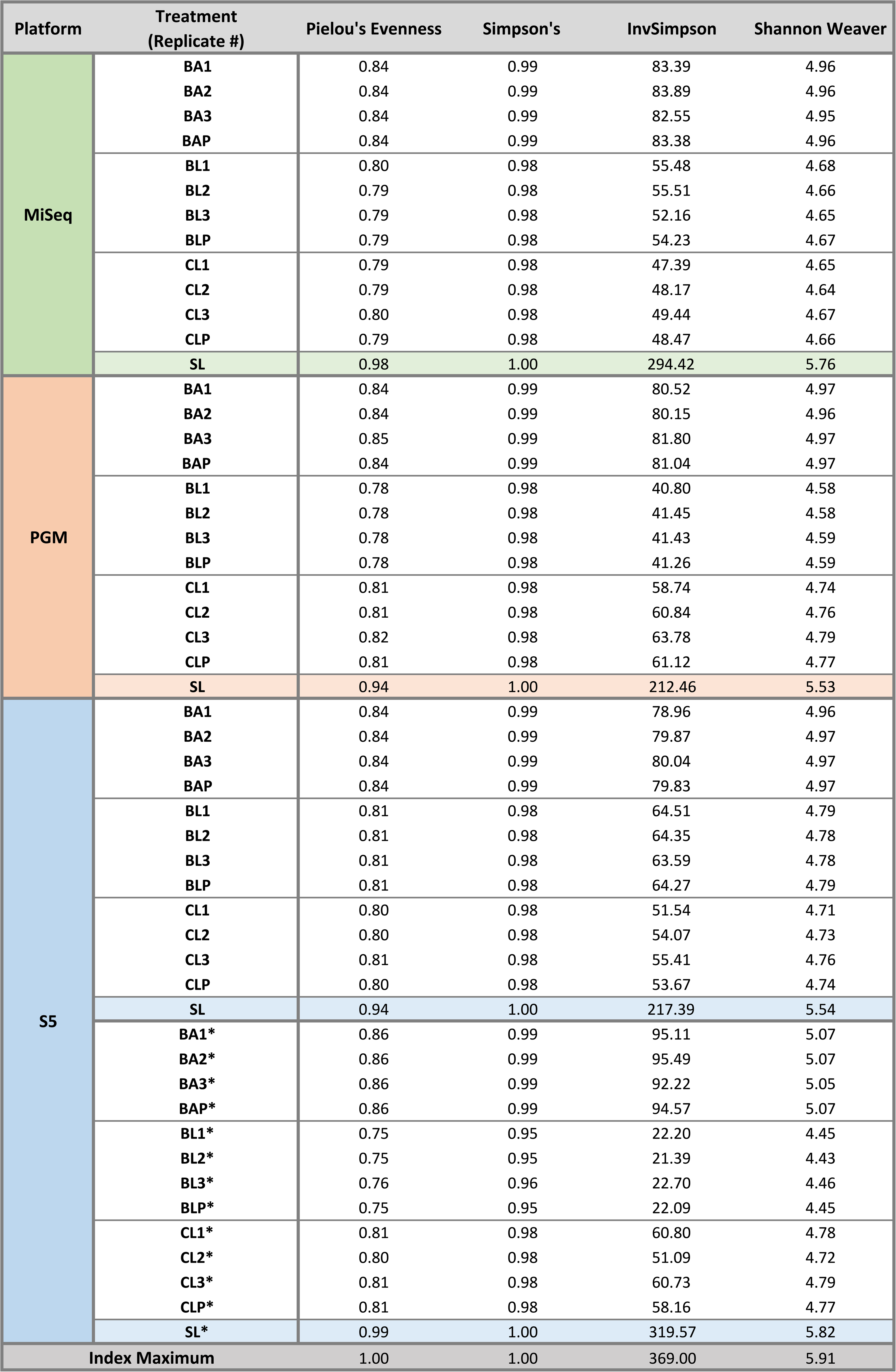
Values for selected diversity indices (Shannon-Weaver, Simpson, Inverse Simpsons, Pielou’s Evenness) for the four amplicon pools (BA = <u>Bulk Abdomen,</u> BL = <u>Bulk Leg,</u> CL = <u>Composite Leg,</u> SL = <u>Single Leg</u>). Replicates are numbered 1–3 while P is the result from pooling the replicates. All results are based on the analysis of a 407 bp amplicon except those marked with a * which are based on a 463 bp amplicon.

### BIN recovery

When the criterion for BIN recovery was set at one or more reads, all platforms recovered >98% of the BINs but only the <u>Single Leg</u> treatment recovered all of them (Figure 4). Differences in recovery success among treatments were greater when the criterion for recovery was set at >0.01% of the reads. Under this criterion, the <u>Single Leg</u> treatment recovered >92.5% of the BINs versus 83–89% for the <u>Bulk Abdomen</u> treatment and about 76–83% for the <u>Composite Leg</u> and <u>Bulk Leg</u> treatments (Table 1). The greater evenness in read count for the <u>Single Leg</u> treatment was striking; it led to lower coefficients of variation, higher diversity indices, and Pielou’s evenness (Table 2; Figure 4; Figure S2). Density plots of read abundance also demonstrated much higher evenness for the <u>Single Leg</u> treatment, especially for the 407 bp amplicon on the MiSeq and for the 463 bp amplicon on the S5 (Figure S3). These differences were also reflected in BIN recovery, Pielou’s evenness, and diversity indices (Table 2; Table S2).

**Figure 3A.**
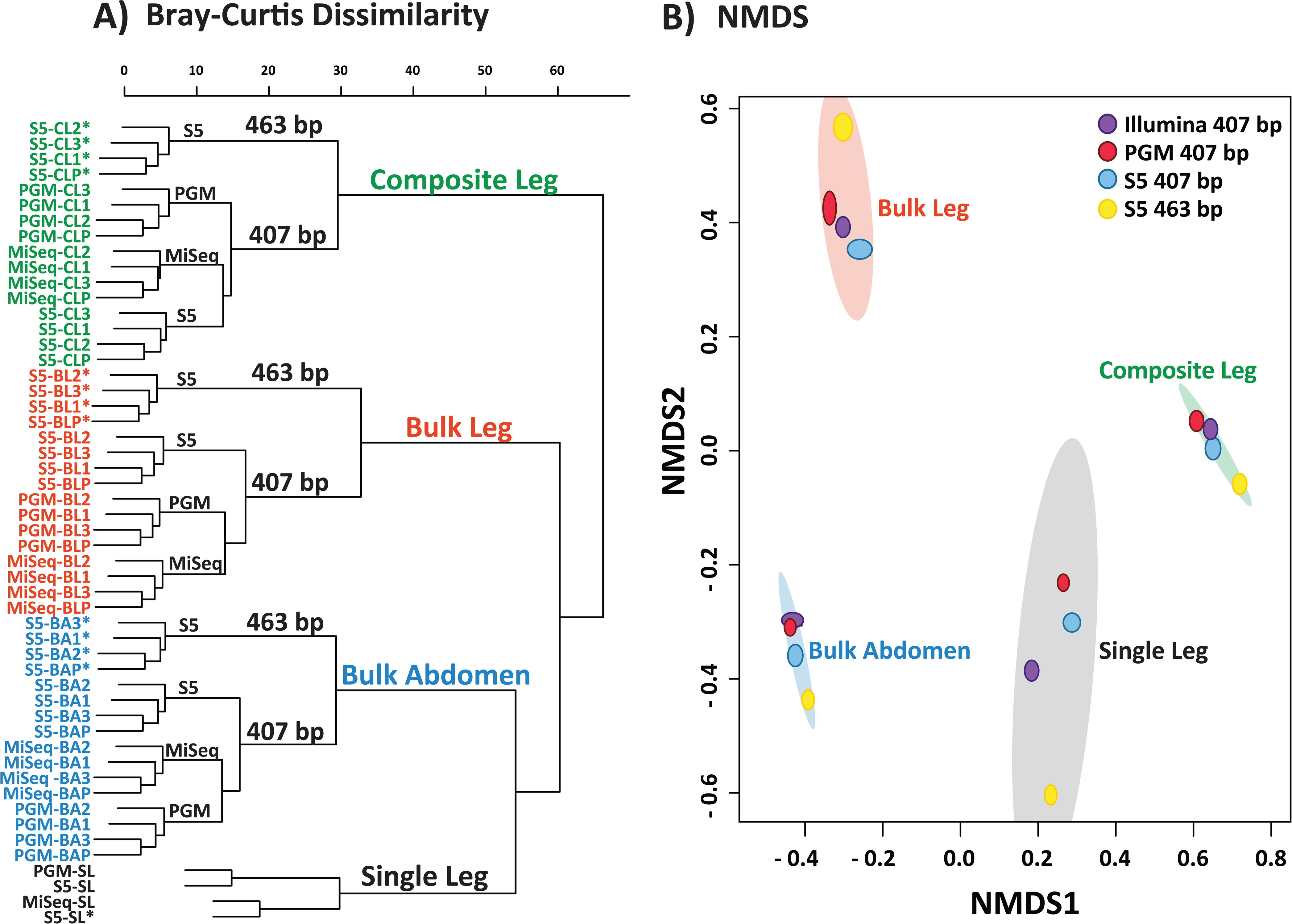
Bray-Curtis Dissimilarity dendrogram for the four amplicon pools (BA = Bulk abdomen, BL = Bulk leg, CL = Composite leg, SL = <u>Single Leg)</u>. Replicates are numbered 1–3 while P is the result from pooling the replicates. The 463 bp amplicon is indicated with an asterisk (*). 3B - Non-metric multidimensional scaling (NMDS) ordinations using Bray-Curtis dissimilarity for the four amplicon pools. Coloured ellipses represent 95% confidence intervals for the BIN composition of the different treatments using ordiellipse (Oksanen et al. 2012). The shapes within each ellipse represent replicates for the four combinations of sequencing platform-amplicon length for three treatments. No replicates were available for the <u>Single Leg</u> treatment, so it has just four points.

**Figure 4.**
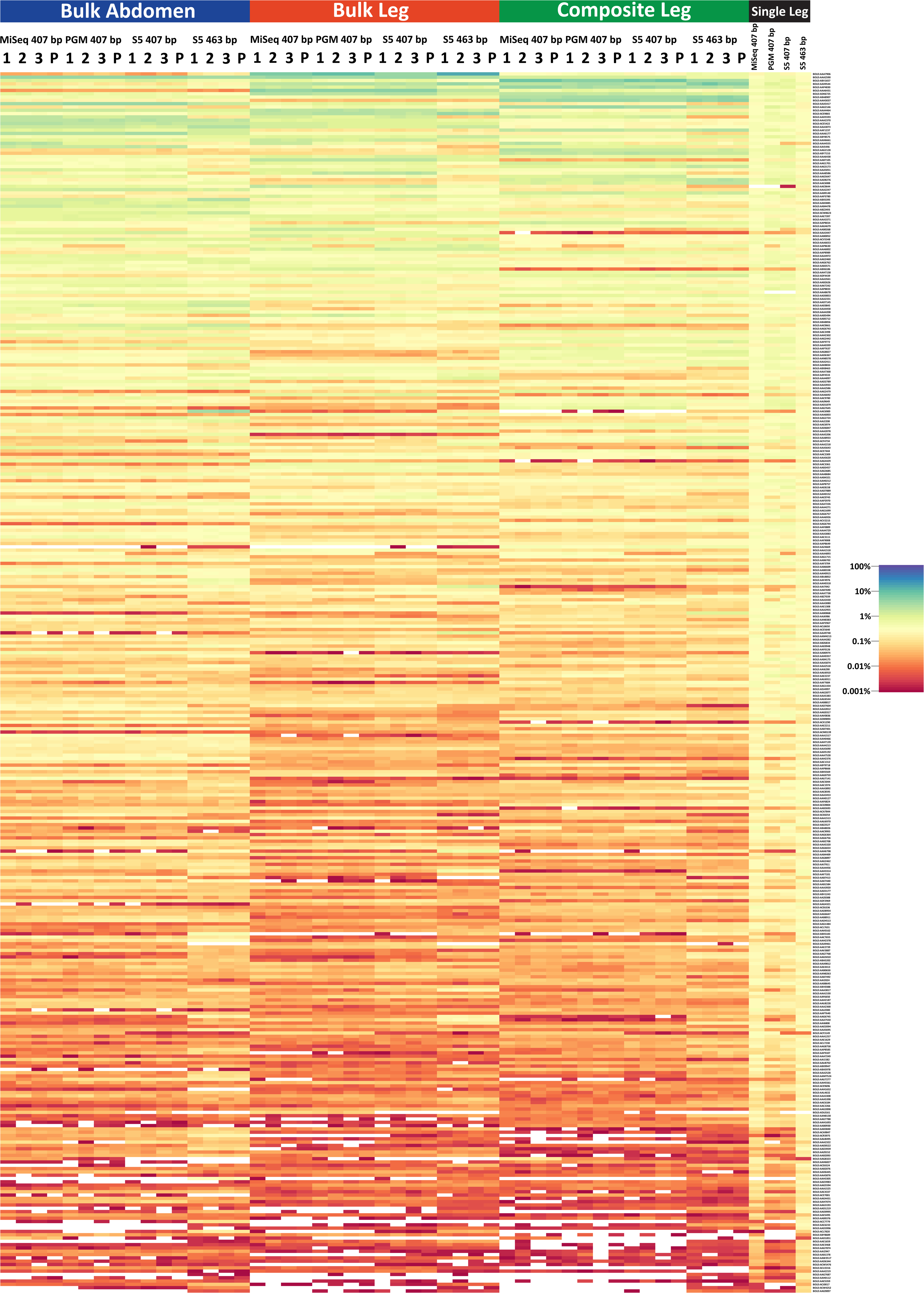
Heat map showing the relative log abundance of the 369 BINs in each treatment for the four amplicon pools. This heat map was created using the JAMP package (https://github.com/VascoElbrecht/JAMP). Technical replicates are indicated with numbers while *in silico* pooled results are designated by the letter P.

### BIN abundances

Because a single specimen of each BIN was included in the mock community, the proportion of sequences from each should, in the absence of bias, be similar across sequencing platforms, amplicons, and treatments. In practice, the relative abundances of the BINs varied markedly. A single-link dendrogram based on Bray-Curtis dissimilarity values indicated that samples clustered first by treatment, next by amplicon length, and finally by sequencing platform (Figure 3A). An analysis of similarity using Bray-Curtis distances affirmed significant differences in BIN abundances by treatment type (p = 0.001, R = 1), amplicon length (p = 0.027, R = 0.17), but not by sequencing platform (p = 0.13, R = 0.037) (Figure 3B; Figure S5).

### Primer mismatches and read count

Examination of the relationship between the read count for each of the 369 BINs and its number of mismatches from the forward primer revealed a strong negative relationship. BINs with a high mismatch count were typically represented by few reads. For example, very few sequences were recovered from the only BIN belonging to the order Dermaptera and this was associated with a high Mismatch Index from the forward primers for the 407 bp and 463 bp amplicons. BIN recovery was substantially higher for the 463 bp amplicon than for the 407 bp amplicon (Table 2; Figure S2). Its superior performance was associated with the fact that its forward primer showed a better match to the DNA extracts. Just 18 BINs had >3 mismatches for the forward primer used to amplify the 463 bp amplicon versus 62 for the forward primer for the 407 bp amplicon (Table S1 and Table S4). The impact of these mismatches was clear; mean read depth and relative abundance of BINs declined after two mismatches for the <u>Bulk Abdomen, Bulk Leg,</u> and <u>Composite Leg</u> treatments and after four mismatches for the <u>Single Leg</u>. When we examined the impact of forward and reverse primer mismatches for a subset of 203 BINs, mean read depth and relative abundance showed a significant decline after four mismatches for the <u>Bulk Abdomen, Bulk Leg,</u> and <u>Composite Leg</u> treatments and after seven for the <u>Single Leg</u>. Kruskal-Wallis tests showed that read depth declined significantly with an increasing number of primer mismatches for the forward primers for both the 463 bp and 407 bp amplicons (p < 0.0001) and for the summed primer mismatches (5’ + 3’) for the subset of 203 BINs (p< 0.0001).

### Impacts of biomass and nucleotide composition on read count

Other factors were also responsible for some of the variation in read counts among BINs. There was, for example, a weak negative correlation between the GC content of an amplicon and its read count, although all values were low (r^2^ < 0.1) excepting the <u>Single Leg</u> treatment on the MiSeq (r^2^ = 0.32). A weak positive correlation (r^2^= 0.24 − 0.28) was also apparent between the abdominal mass of a BIN and its read count on all platforms.

### Non-Target Sequences

Every run recovered some sequences with substantial sequence divergence from the Sanger reference library (Table 1). The incidence of these non-target sequences for the 407 bp amplicon was slightly lower (4 – 6%) on the PGM and S5 platforms than on the MiSeq (8 – 10%). Interestingly, the 463 bp amplicon had substantially more non-target reads (15 – 17%). Many of the non-target reads were chimeras (8 – 81 %; Table 1). After their exclusion, most sequences assigned to an OTU did not find a match to a sequence in the supplemental libraries. Of those that did, similar numbers matched to a known bacterial sequence or to another arthropod.

### Taxonomic Bias

There was also evidence of taxonomic bias in the read counts for BINs between the two amplicons. For example, Orthoptera, Lepidoptera, and Diptera dominated the 407 bp sequences from the <u>Bulk Abdomen</u> and <u>Bulk Leg</u> treatments while Lepidoptera, Mecoptera, Diptera, and Coleoptera dominated the 463 bp amplicon (Table S5). The 463 bp amplicon also showed more variation among treatments than the 407 bp amplicon (Table S5). Few sequences were recovered for Dermaptera, especially for the 407 bp amplicon, likely reflecting its possession of 5 mismatches from the forward primer (Table S1). Among the bulk samples, relative abundance differed among treatments. For example, the relative abundance of Lepidoptera and Mecoptera was lower, while Diptera and Orthoptera were higher in the <u>Composite Leg</u> than in the <u>Bulk Leg</u> and <u>Bulk Abdomen</u> treatments. The proportion of read counts for Trichoptera showed particularly large variation, being 5–25X higher for the <u>Bulk Leg</u> than the <u>Bulk Abdomen</u> and <u>Composite Leg</u> treatments across all platforms and for both amplicons.

## Discussion

Metabarcoding is a powerful tool for characterizing biodiversity patterns (Cristescu 2014), but data interpretation is complicated by several factors. PCR amplification bias and variation in the copy number of template DNA from the source specimens not only make it impossible to estimate abundances, but can impede the recovery of all species (Yu et al. 2012; Li et al. 2013; Beng et al. 2016). Although prior studies have revealed these complexities, there has been limited evaluation of the strength of their influence on interpretations of taxon diversity. To address this gap, the present study has examined the impacts of diverse factors including source DNA, PCR primers, sequencing platform, and sequencing depth on species recovery from a mock community of arthropods.

### Sequencing Depth

Variation in sequencing depth (read count) can directly impact taxon recovery and hence perceived diversity patterns. This is particularly true for comparisons among datasets with differing read counts (Leray et al. 2015; Leray and Knowlton 2017; Elbrecht et al. 2017). Low read depth typically means that some rare species with little biomass in a community will be overlooked, leading to underestimation of alpha diversity. When taxon counts are incomplete, comparisons among sites are also compromised (Bellemain et al. 2012; Sickle et al. 2015; Yamamoto et al. 2017), producing overestimates of beta diversity (Sickle et al. 2015). Both rarefaction and species accumulation curves are valuable for ascertaining if sequencing depth has been adequate. When sequences have been recovered from all species, the slope of the rarefaction curve is zero, providing a simple criterion for gauging the adequacy of read coverage (Lanzen et al. 2017). In real world situations, the true species count is unknown and increased sampling effort nearly always raises the species count, meaning there is no asymptote. Although taxon richness was fixed in our study, we employed a slope of 0.01 as the criterion for assessing when taxon diversity had achieved an asymptote as this approach can be employed in studies on natural communities. Under this criterion, there were substantial differences among the four treatments and between primer sets.

Analysis of BIN accumulation curves indicated that read depth was sufficient for all four treatments to achieve a slope of 0.01. However, the <u>Single Leg</u> treatment reached this value with much lower read depth than the bulk samples due to its relative protection from the impacts of PCR bias (Nichols et al. 2017; Pan et al. 2014, Dabney and Meyer 2012; Elbrecht and Leese 2015). Interestingly, the other three treatments showed similar BIN accumulation curves on all three sequence platforms, suggesting shared factors are constraining BIN recovery.

### Sequencing Platforms

The three sequencing platforms generated similar estimates of BIN diversity. However, results from the MiSeq had advantages over those from the PGM and S5. Its paired end protocol consistently recovered sequences for the full 407 bp amplicon, which can be a requirement for clustering algorithms and is also useful for haplotype analysis (Elbrecht et al. 2018), while those from the other platforms were often truncated. Its reads also possessed fewer indels than those from PGM and S5. Finally, the MiSeq reads had consistently higher QV across the amplicon. Because these factors simplified data analysis (Mardis et al. 2013; Edgar et al. 2013) and sequencing costs were similar, the MiSeq is currently the best platform for metabarcoding (Mardis 2013). However, because it cannot analyze amplicons longer than 500 bp, third-generation sequencing platforms (Tedersoo et al. 2018; Hebert et al. 2018; Wilkinson et al. 2017) will be an attractive option if their current limitations on read number (Pacific Biosciences) and quality (Oxford Nanopore) are overcome.

### Impacts of Analytical Protocols

Our four treatments made it possible to compare the impact of targeting different tissues, employing different DNA extraction regimes, and using different PCR protocols. Despite their similar tissue input and DNA extraction regime, the <u>Single Leg</u> treatment achieved asymptotic diversity much more rapidly than the <u>Composite Leg</u> treatment, indicating how separate PCR reactions reduce amplification bias. By contrast, BIN accumulation curves for the <u>Composite Leg</u> treatment were similar to those for the <u>Bulk Leg</u> and <u>Bulk Abdomen,</u> indicating that DNA extraction was equally effective whether carried out on single specimens or on bulk samples. Comparison of the results for the bulk/composite samples did reveal more nuanced differences as they showed the number of reads for particular taxa varied among these three treatments despite similar BIN recovery profiles. These differences likely stem from differential leg/abdomen mass ratios among species which varied the mitochondrial copy number for the component species among treatments. Certainly, mitochondrial copy number varies among tissues and among species (Veltri et al. 1990; Cole 2016). Future efforts to explore this relationship and its importance to metabarcoding studies must quantify copy number differences between tissues and species. In the absence of such information, copy number bias can be reduced by partitioning the specimens in a bulk sample into a few size fractions (Elbrecht et al. 2017A; Vivien et al. 2016).

Variation in read counts for the taxa in any bulk sample are also influenced by primer bias. In this study, these effects were indicated by the linkage between the read counts for each BIN and the number of mismatches between its COI sequence and the primer set. Although degenerate primers (Yu et al. 2012; Elbrecht and Leese 2017; Moriniere et al. 2016) and improved primer sets (Clarke et al. 2014; Leray and Knowlton 2017; Elbrecht and Leese 2017) can reduce such bias, they cannot conquer the problem unless all target species possess identical sequences for the primer binding sites, a condition that will never be satisfied for a large assemblage. However, efforts to target highly conserved regions can improve the situation. For example, the BIN accumulation curve reached its asymptote with much lower read coverage for the 463 bp than the 407 bp amplicon. More effort is needed to develop better primer sets by testing their performance on a breadth of taxa and to minimize mismatches by sorting samples into taxonomic groups (Moriniere et al. 2016; Bellemaine et al. 2012; Cristescu 2014; Tedersoo et al. 2015). Efforts to minimize specimen bias should include mass and taxonomic sorting to reduce differences in template DNA quantity between specimens (Elbrecht et al. 2017A; Moriniere et al. 2016; Vivien et al. 2016). Currently, the only means to circumvent biases is to process specimens individually (e.g. <u>Single Leg</u> treatment), which is so time consuming and costly that it is difficult to implement for large biodiversity surveys (Ji. et al 2013).

### BIN recovery

Most of the 369 BINs in the template pools were recovered in all four treatments. However, this outcome shifts if recovery success is defined as those BINs comprising at least 0.01% of the read count, a criterion often employed to minimize the impacts of sequencing errors, chimeras, and contaminants (Leray and Knowlton 2017). Under this criterion, BIN recovery was high (> 92.5%) for the <u>Single Leg</u> treatment, but substantially lower (76% - 89%) for the other three. Interestingly, the <u>Bulk Abdomen</u> treatment showed higher BIN recovery than the <u>Bulk Leg</u> and <u>Composite Leg</u> treatments, perhaps reflecting more similar mitochondrial copy numbers among abdomens than legs (Veltri et al. 1990; Cole 2016). As expected, BIN recovery was higher for the 463 bp than the 407 bp amplicon.

### False positives and negatives

Up to 26 of all BINs failed to achieve a 0.01% read abundance in the bulk and composite treatments meaning they would often be excluded during analysis, creating false negatives that would underestimate alpha diversity. As in other metabarcoding studies (Vivien et al. 2016; Ficetola et al 2014; Brandon-Mong et al. 2015; Port et al. 2015), false positives were also encountered, likely reflecting eDNA associated with specimens, contamination during sample processing (Port et al. 2015) or NUMTs (Song et al. 2008). Their impact can be reduced by employing curated reference libraries to both recognize sequences derived from known species and to exclude paralogs and pseudogenes (Hebert et al. 2003; Landi et al. 2014; Zimmerman et al. 2014; Braukmann et al. 2017; Bergsten et al. 2014). In addition, the use of negative controls is an effective way to evaluate the incidence of contaminants introduced during sample processing (Port et al. 2015).

### Conclusions

This study has examined the complexities encountered in evaluating the species composition of a mock community comprised of 374 species of arthropods. Some results were reassuring. Similar measures of overall taxon diversity were obtained from different sequencing platforms, from different tissues, from different DNA extraction protocols, and from different PCR primers. However, this congruence needs to be qualified. Firstly, this study has shown that the analytical effort required to obtain comprehensive information on species composition through the analysis of bulk samples is far higher than that required to obtain the same information through specimen-based protocols. For example, the Sanger sequencing of 369 specimens would deliver precise information on the species composition and abundance of each sample. By comparison, the recovery of a complete species list by sequencing a merged pool of amplicons following separate extraction and PCR (e.g. <u>Single Leg</u> treatment) required 60,000 reads. When samples were pooled prior to PCR, the recovery of a near-complete species list required at least 250,000 reads, and the proportion of the taxa recovered were represented by so few reads (<0.01%) that they could be excluded during data cleansing. It is worth emphasizing that many natural communities present greater analytical complexity than the mock assemblage examined in this study – they include more species and the abundances of these species show great variation. Given these complications, it is clear that community characterization through metabarcoding will often require both intensive sequencing effort and improved approaches to discriminate between those low frequency reads that are spurious and those that derive from rare species.

## Acknowledgements

We thank the lab and collections staff at the Centre for Biodiversity Genomics for aiding in acquiring and processing the specimens in this study and Suzanne Bateson for aid with graphics. We are also grateful to Jenna Quinn and other staff at the rare Charitable Research Reserve for facilitating the collection of specimens. This study was enabled by support from the Ontario Ministry of Research, Innovation and Science and from the Canada First Research Excellence Fund to the ‘Food From Thought’ research program.

---

Figure S1A – Phred or quality (QV) scores across read length for 407 bp amplicon on the Illumina MiSeq, Ion Torrent PGM, and Ion Torrent S5 as well as for 463 bp amplicon on the Ion Torrent S5. S1B – Histogram of read lengths for the three platforms and two amplicon lengths on the Ion Torrent S5.

Figure S2 – Renyi’s diversity graphs using the pooled replicates for the <u>Bulk Abdomen, Bulk Leg,</u> and <u>Composite Leg</u> treatments and the single replicate for the <u>Single Leg</u> treatment.

Figure S3 – Density plots based on the relative abundance for the 369 BINs common across all treatments. The results from <u>Bulk Abdomen, Bulk Leg, Composite Leg,</u> and <u>Single Leg</u> PCR are represented by blue, green, red, and black lines respectively. Each panel represents a different sequencing platform: A) 463 bp on the Ion Torrent S5, B) 407 bp on the Ion Torrent S5, C) 407 bp on the Illumina MiSeq, and D) 407 bp on the Ion Torrent PGM.

Figure S4 – Jaccard Similarity dendrogram for the four amplicon pools (BA = <u>Bulk Abdomen,</u> BL = <u>Bulk Leg,</u> CL = <u>Composite Leg,</u> SL = <u>Single Leg)</u>. Replicates are numbered 1–3 while P is the result from pooling the replicates. The 463 bp amplicon is indicated with an asterisk (*).

Figure S5 – Non-metric multidimensional scaling (NMDS) ordinations using Bray-Curtis dissimilarity for the four amplicon pools (BA = <u>Bulk Abdomen,</u> BL = <u>Bulk Leg,</u> CL = <u>Composite Leg,</u> SL = <u>Single Leg)</u>. Coloured ellipses represent 95% confidence intervals for the BIN composition of the different treatments using ordiellipse (Oksanen et al. 2012). No replicates were available for the <u>Single Leg</u> treatment, so it has just four points.

Table S1 – Taxonomy and results for the mock communities used in this study. A) includes the read depth for each BIN while B) shows the relative abundance of each BIN in the different mock communities for the four amplicon pools (BA = <u>Bulk Abdomen,</u> BL = <u>Bulk Leg,</u> CL = <u>Composite Leg,</u> SL = <u>Single Leg</u>). Replicates are numbered 1–3 while P is the result from pooling the replicates. The 463 bp amplicon is indicated with an asterisk (*).

Table S2 – Primer sequences for the different platforms. A) primer sequences for Ion Torrent PGM and S5 and B) primers for the Illumina MiSeq including the COI primer (red) and unique molecular identifiers (UMIs; green). The Nextera Transposase adapters have two components: an adaptor sequence (yellow) and a sequencing primer (purple). Sequencing adaptors (Ion Torrent PGM and S5) and flow cell adapters (Illumina) are shown in blue.

Table S3 – Slopes from rarefaction curve at the prior 1, 2, 3, 5, and 10 points for the four amplicon pools (BA = <u>Bulk Abdomen,</u> BL = <u>Bulk Leg,</u> CL = <u>Composite Leg,</u> SL = <u>Single Leg)</u>. Replicates are numbered 1–3 while P is the result from pooling the replicates. The 463 bp amplicon is indicated with an asterisk (*).

Table S4 – Summary table of primer mismatches for the 463 bp and 407 bp amplicons for A) each BIN and B) mean read depth per primer mismatch for the four amplicon pools (BA = <u>Bulk Abdomen,</u> BL = <u>Bulk Leg,</u> CL = <u>Composite Leg,</u> SL = <u>Single Leg</u>). Replicates are numbered 1–3 while P is the result from pooling the replicates. The 463 bp amplicon is indicated with an asterisk (*).

Tables S5 – Read depth and relative abundance for each treatment and replicate by A) Order and B) Family for the four amplicon pools (BA = <u>Bulk Abdomen,</u> BL = <u>Bulk Leg,</u> CL = <u>Composite Leg,</u> SL = <u>Single Leg</u>). Replicates are numbered 1–3 while P is the result from pooling the replicates. The 463 bp amplicon is indicated with an asterisk (*).

## Other Supplementary material

MBRAVE_OTUmerger_MER.R – R script for combining mBRAVE OTU tables into single file.

OTU_subsampler_MER.R – R Script for generating rarefraction curves from an OTU table.

